# *In vitro* screening of herbal medicinal products for their supportive curing potential in the context of SARS-CoV-2

**DOI:** 10.1101/2021.03.01.433344

**Authors:** Hoai Thi Thu Tran, Philipp Peterburs, Jan Seibel, D. Abramov-Sommariva, Evelyn Lamy

## Abstract

**Background:** Herbal medicinal products have a long-standing history of use in the therapy of common respiratory infections. In the COVID-19 pandemic, they may have the potential for symptom relief in non-severe or moderate disease cases. Here we describe the results derived by *in vitro* screening of five herbal medicinal products with regard to their potential to i) interfere with the binding of the human Angiotensin-converting enzyme 2 (ACE2) receptor with the SARS-CoV-2 Spike S1 protein, ii) modulate the release of the human defensin HBD1 and cathelicidin LL-37 from human A549 lung cells upon Spike S1 protein stimulation and iii) modulate the release of IFN-γ from activated human peripheral blood mononuclear cells (PBMC). The investigated extracts were: Sinupret extract (SINx), Bronchipret thyme-ivy (BRO TE), Bronchipret thyme-primrose (BRO TP), Imupret (IMU), and Tonsipret (TOP).

**Methods:** The inhibitory effect of the herbal medicinal products on the binding interaction of Spike S1 protein and the human ACE2 receptor was measured by ELISA. The effects on intracellular IFN-γ expression in stimulated human PBMCs were measured by flow cytometry. Regulation on HBD1 and LL-37 expression and secretion was assessed in 25d long-term cultured human lung A549 epithelial cells by RT-PCR and ELISA.

**Results:** IMU and BRO TE concentration-dependently inhibited the interaction between spike protein and the ACE2 Receptor. However, this effect was only observed in the cell-free assay at a concentration range which was later on determined as cytotoxic to human PBMC. SINx, TOP and BRO TP significantly upregulated the intracellular expression of antiviral IFNγ from stimulated PBMC. Co-treatment of A549 cells with IMU or BRO TP together with SARS-CoV-2 spike protein significantly upregulated mRNA expression (IMU) and release (IMU and BRO TP) of HBD1 and LL-37 (BRO TP).

**Conclusions:** The *in vitro* screening results provide first evidence for an immune activating potential of some of the tested herbal medicinal extracts in the context of SARS-CoV-2. Whether these could be helpful in prevention of SARS-CoV-2 invasion or supportive in recovery from SARS-CoV-2 infection needs deeper understanding of the observations.

## Introduction

All over the world the population is struggling with the outbreak of COVID-19. Effective antiviral therapies are unavailable so far, comprehensive vaccination has yet to be achieved and various mutants of the enveloped, single-stranded RNA virus SARS-CoV2 have emerged (Greaney et al., 2021). Thus it is of critical importance to search for compounds, which could help in fighting the virus pandemic. Testing the effectiveness of different drug types used previously in the treatment of other diseases is here one strategy to accelerate the development. Herbal medicinal products have the potential to interfere with various steps of the viral replication cycle (Brendler and Al-Harrasi, 2020; Mani et al., 2020). Besides, they have been reported to exhibit an anti-inflammatory and immune modulating potential and thus may also be supportive in prevention or attenuation of mild to moderate SARS-CoV-2 infections (Glatthaar-Saalmüller B, 2011; Seibel et al., 2015; Seibel et al., 2018). In this pilot screening study, five herbal medicinal products, marketed for the treatment against respiratory infections (Bachert, 2020; Rossi et al., 2012; Seibel et al., 2015; Seibel et al., 2018), were explored for their potential to i) interfere with the binding of the human Angiotensin-converting enzyme 2 (ACE2) receptor with the SARS-CoV-2 spike protein. The ACE2 receptor found in various organs including type I and II pneumocytes, endothelial cells, oral and nasal mucosa but also intestinal tissues, liver, kidney, or brain (Hamming et al., 2004; Xu et al., 2020) has been identified as the key cellular receptor, which facilitates uptake of the SARS-CoV-2 virus into the host cell (Ortiz et al., 2020). Drugs targeting the interaction between the spike protein receptor binding domain of SARS-CoV-2 and ACE2 receptor may thus offer some protection against this novel viral infection (Hu et al., 2020) (Yang et al., 2020). ii) We investigated whether the extracts can modulate the release of the defensin HBD1 and cathelicidin LL-37 from human A549 lung cells upon SARS-CoV-2 spike protein stimulation. In the innate immune system, these antimicrobial peptides (AMPs), have a non-enzymatic inhibitory effect on a broad spectrum of microorganisms (Wilson et al., 2013). Human beta defensins (HBDs) have known antiviral effects on both enveloped and non-enveloped viruses (Ding et al., 2009). Due to the relative nonspecificity of the targets of defensins compared to those of the adaptive arm, antiviral applications of defensins are conceptually ideal for protection against different viral infections (Holly et al., 2017; Wilson et al., 2013). iii) We investigated whether the plant extracts could modulate the activated immune system in terms of regulating anti-viral interferon gamma (IFN-γ) production. This type II interferon is produced by T lymphocytes and NK cells and essential for antiviral defense. It suppresses virus replication and activates T cell cytokine production (Levy and García-Sastre, 2001).

## Methods

### Extracts

All plant extract mixtures were provided by Bionorica SE either in dried or fluid form. The dried extracts were used in different dilutions. The following extracts (Bionorica SE, Neumarkt in der Oberpfalz, Germany) were investigated:

Bronchipret® thyme-ivy (BRO TE), an extract of thyme herb (*Thymus vulgaris* L. or *Thymus zygis* L.) and ivy leaves (*Hedera helix* L.). BRO-TE is a mixture of fluid extracts of thyme herb (extractant: ammonia solution 10% (m/m) / glycerol (85%) (m/m) / ethanol 90% (v/v) / water (1:20:70:109); drug–extract ratio (DER): 1:2–2.5) and ivy leaves (extractant: ethanol 70% (v/v); DER: 1:1) as contained in Bronchipret® syrup with a thyme/ivy fluid extract ratio of 10:1. In order to minimise ethanol content in the test system the extract mixture was dealcoholised by rotary evaporation to a final ethanol content of 1% (v/v). To control for loss of volatile ingredients, specific identity tests were performed with the concentrate.

Bronchipret® thyme-primrose (BRO TP), an extract of thyme herb (*Thymus vulgaris* L. or *Thymus zygis* L.) and primrose root (*Primula veris* L. or *Primula elatior* (L.) Hill). BRO-TP is a mixture of genuine dry extracts of thyme herb (extraction solvent: ethanol 70% (v/v); DER: 6–10:1) and primrose root (extractant: ethanol 47% (v/v); DER 6–7:1) as contained in Bronchipret® TP film-coated tablets without excipients and with a final thyme/primrose dry extract ratio of 2.67:1. A stock solution of 100 mg/ml was prepared in 50% EtOH

Imupret® (IMU), 100 g Imupret oral drops contain: 29 g of an ethanolic-aequous extract (extraction solvent: Ethanol 59 Vol.-%) out of Marshmellow root *(Altheae officinalis* L.) 0.4 g, Chamomille flowers (*Matricaria recutita* L.) 0.3 g, Horsetail herb (Equisetum avense L.) 0.5 g, Walnut leafs (*Juglans regia* L.) 0.4 g, Yarrow herb (*Achillea millefolium* L.) 0.4 g, Oak bark (*Quercus robur* L.) 0.2 g, Dandelion herb (*Taraxacum officinale* F.H. Wiggers) 0.4 g. Total-ethanol 19 % (v/v)‥ In order to minimise ethanol content in the test system the extract mixture was dealcoholised (> 0.5% (v/v)) by rotary evaporation. The content quality of the dealcoholized test item complied with Imupret® oral drops as checked by identity tests and quantitative analysis.

Sinupret® extract (SINx), combined genuine dry extract (BNO 1011) of gentian root (*Gentiana lutea* L.), primrose flower (*Primula veris* L.), sorrel herb (*Rumex crispus* L.), elder flower (*Sambucus nigra* L.) and verbena herb (*Verbena officinalis* L.) with a ratio of 1:3:3:3:3 (extraction solvent: ethanol 51% (v/v); DER 3–6:1) as contained in Sinupret® extract coated tablets without excipients.

Tonsipret® (TOP), homeopathic dilution for tonsillitis tablets containing 37.5% Dilution Capsicum D3 (*Capsicum annuum* L.), 37.5% Dilution Guajacum D3 (*Guaiacum officinale* L./ *Guaiacum sanctum* L.) and 25.0% mother tincture Phytolacca (*Phytolacca americana* L.). In order to minimise ethanol content in the test system the mixture was dealcoholized (> 0.5% (v/v)) by rotary evaporation. The quality of the dealcoholised test item complied with the corresponding manufacturing stage of the herbal medicinal product Tonsipret® as checked by identity analyses. All extracts were centrifuged (16.000 x g, 3 min, RT) and the supernatant was used for the experiments. The final concentration of the solvent (ethanol) was less than 0.5% in all assays.

### Human A549 lung cell line and primary human PBMC

The human lung adenocarcinoma A549 cell line was purchased from DSMZ (Germany). For the experiments, cells were used at low passage number after thawing. Cells were cultured in RPMI medium containing 10% heat inactivated FCS, 1% penicillin/streptomycin and 1% L-glutamine for 25 days for differentiation according to the protocol by Cooper et al., 2016(Cooper et al., 2016). After 25 days, cells were exposed to the extracts for different time points either with or without co-treatment with SARS-CoV-2 spike protein (Trenzyme, Germany).

For PBMC isolation, blood was taken in Li-Heparin vacutainers from healthy volunteers at the University of Freiburg - Medical Center after written informed consent. The study was approved by the Ethics Committee of the University of Freiburg and carried out according to their guidelines and regulations (ethical vote 373/20). PBMC isolation was done by centrifugation on a LymphoPrepTM gradient using SepMate centrifugation tubes (Stemcell Technologies, Germany), cells were then washed twice with PBS and viability and concentration was determined using the trypan blue exclusion test.

### Assessment of SARS-CoV-2 Spike-ACE2 binding inhibition

The capacity of extracts to inhibit the SARS-CoV-2-Spike-ACE2 binding was tested using the SARS-CoV-2 (COVID-19) Inhibitor screening kit (BioCat GmbH, Heidelberg, Germany) according to the manufacturer’s instructions. This colorimetric ELISA assay measures the binding between immobilized SARS-CoV-2 Spike protein RBD and biotinylated human ACE2 protein. The colorimetric detection is done using streptavidin-HRP followed by TMB incubation.

### Quantification of human defensins by qRT-PCR

A549 cells were treated with extracts with/without Sars-CoV-2 Spike S1 protein (Trenzyme, Germany) for different time points. Total RNA was isolated from the cells using the RNeasy mini isolation kit from Qiagen (Germany) with a purification step using the RNase-free DNase kit (Qiagen) according to the manufacturer’s instructions. The quantity and quality of RNA were determined by a NanoDrop ND-1000 spectrophotometer (Thermo Scientific, Germany). 25 ng RNA was used for qPCR reaction. Changes in mRNA levels were measured by qRT-PCR as described previously (Nam et al., 2006; Pierson et al., 2013).

### Quantification of HBD1 and LL-37 peptide release from A549 cells

HBD1 and LL-37 peptide release was quantified in supernatants from 25 day long-term cultured A549 cells using a human BD-1 Standard ABTS ELISA Development Kit (PeproTech, Germany) and LL-37 human ELISA kit (HycultBiotech, Germany) according to the manufacturer’s instructions.

### Flow cytometric analysis of intracellular IFN-gamma expression

PBMC were treated with extracts for 6h and co-stimulated with 50 ng/ml phorbol 12-myristate 13-acetate (PMA), 1 μg/ml ionomycin, and a SARS-CoV-2 S peptide pool (Miltenyi Biotec, Germany) for immune reaction together with 10 μg/ml brefeldin A. Cells were then collected and processed for intracellular staining with anti-IFN-γ-FITC monoclonal antibody (Miltenyi Biotec, Germany). Changes in the subset of lymphocyte cells and cytokine production (IFN-γ) were assessed using flow cytometry (FACS Calibur, BD, Germany).

### Determination of cell viability

The LIVE/DEAD Fixable Far Red Dead Cell Stain Kit (Thermo Fisher Scientific, Germany) and the LDH-Glo Cytotoxicity Assay (Promega, Germany) were used to determine cytotoxicity of extracts in PBMC and A549 cells, respectively after 24 hour exposure, according to the manufacturer’s instructions.

### Statistics

Results were analysed using the GraphPad Prism 6.0 software (La Jolla, California, USA). Data were presented as means + SD. Statistical significance was determined by the one way ANOVA test followed by Holm-Sidak’s correction. P values <0.05 (*) were considered statistically significant and <0.01 (**) were considered highly statistically significant.

## Results

### Effect of extracts on SarsCoV2-Spike-ACE2 binding inhibition

The inhibitory potential of the extracts between the SARS-CoV-2 spike protein and the ACE2 receptor was tested in a cell free assay. A concentration-dependent inhibitory effect on spike binding to ACE2 was seen with extract IMU (maximum of 86% ± 1.3), extract BRO TP (maximum of 44% ± 16.8) and extract BRO TE (maximum of 77% ± 10.8). For extracts TOP, no relevant effect (data not shown) and for SINx minor effects were observed (figure 1).

**Figure 1:**
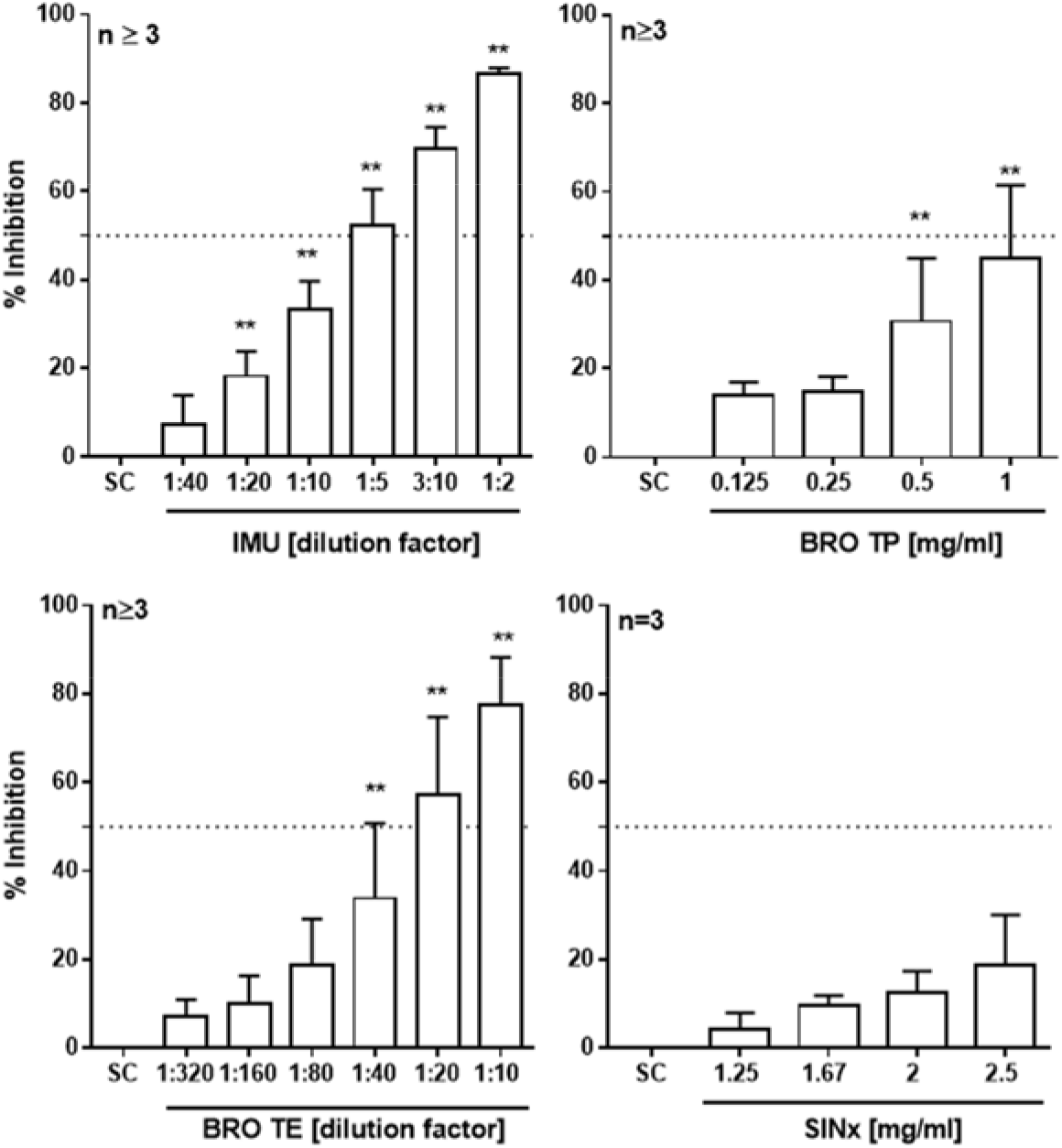
Efficacy of Spike-ACE2 binding inhibition by the tested extracts. The SARS-CoV-2 S protein RBD coated plate was incubated with plant extracts and biotinylated human ACE2 for 1 h. Thereafter, the plate was washed thoroughly and incubated with Streptavidin-HRP followed by colorimetric detection using a multiplate reader from Tecan (Germany). Bars are means +SD of at least three independent experiments.

### Effect of extracts on IFN-γ intracellular expression in activated human PBMC

First, cytotoxicity of the extracts was quantified after 24 h exposure in human PBMC. Relevant toxicity (i. e. >10%) was evident for extract IMU at ≥ 1:40 dilution, for extract BRO-TP at ≥ 0.25 mg/ml and for BRO-TE at ≥ 1:200 dilution. For TOP and SINx, no relevant toxicity could be seen in the applied *in vitro* model. Upon stimulation with a mixture of SARS-CoV-2 S peptide pool, PMA and ionomycin, a significant increase in intracellular IFN-γ expression (in range of 7-15%) could be seen (see figure 2). Compared to control cells, a significant increase in IFN-γ expression could be seen in stimulated PBMC with extract TOP (145% ± 27 at a 1:60 dilution), extract SINx (169% ± 26 at 1.25 mg/ml) and extract BRO TE (157% ± 31 at a 1:1000 dilution) (figure 2). For extract IMU and BRO TP, no additional increase in IFN-γ expression was seen. In contrast, at ≥ 0.167 mg/ml, extract BRO TP had an inhibitory effect on INF-γ, which is likely due to beginning cytotoxicity.

**Figure 2:**
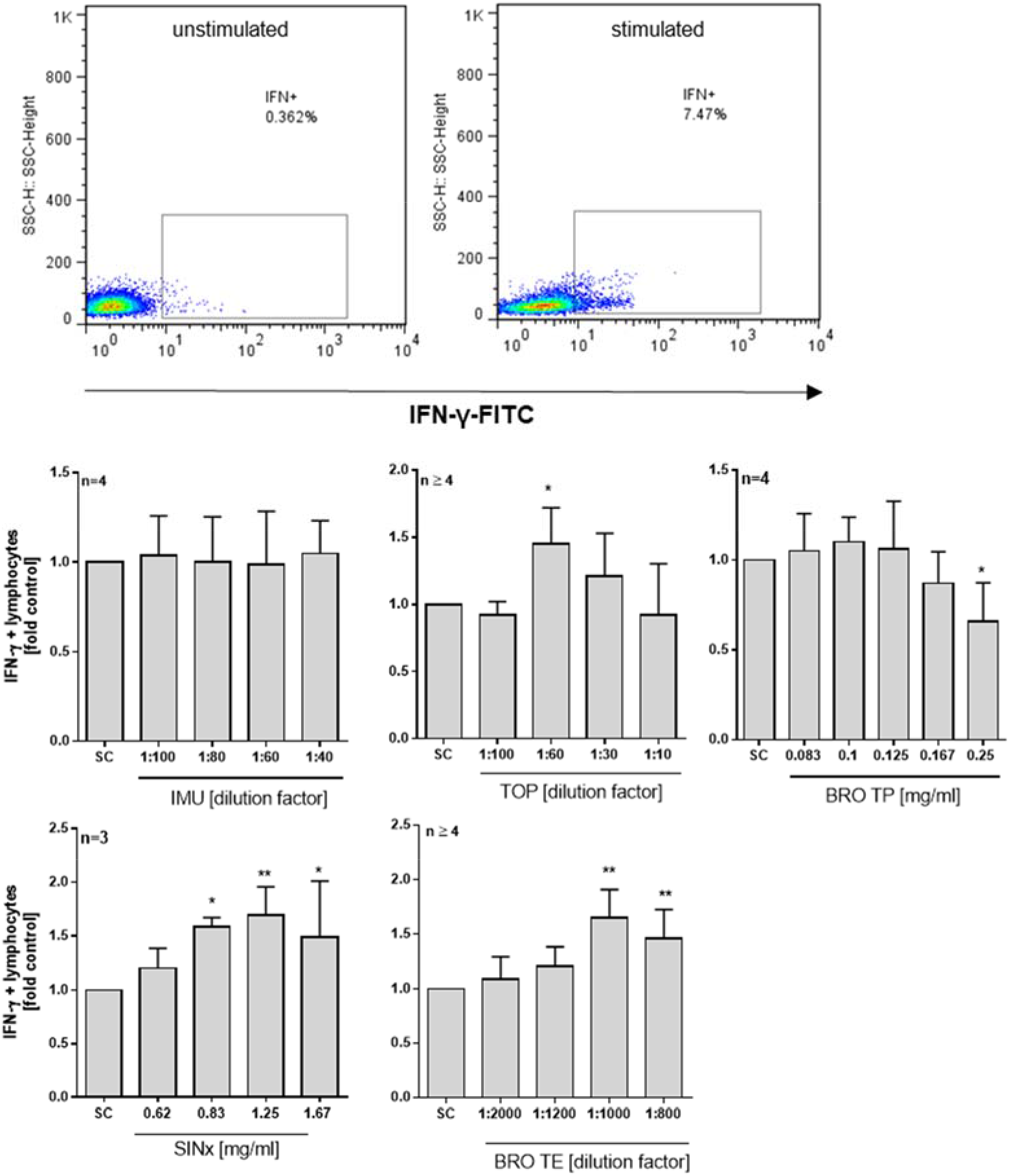
Intracellular IFN-γ expression from stimulated human PBMC upon extract treatment. Cells were stimulated with a mixture of SARS-CoV-2 S peptide pool, PMA and ionomycin and treated with the extracts for 6 hours. Brefeldin A was used to enhance intracellular cytokine staining signals. Representative scattergrams of intracellular INF-γ staining without (left) and after (right) stimulation are given. Bars are means +SD of at least three independent experiments.

### Effects of extracts on mRNA expression and peptide release of HBD1and LL-37

For first insights into the effects of the test items on cellular defense mechanisms we analysed the mRNA expression and secretion of HBD-1 and LL-37 in human A549 cells. Based on the findings in human PBMC, the absence of relevant cytotoxicity (i. e. > 10%) upon extract exposure was first confirmed in 25d long-term cultured A549 cells using the LDH-Glo cytotoxicity assay (data not shown). We then treated the cells with Sars-CoV-2 Spike S1 protein together with the extracts for 24 h. Thereafter, a significant increase in HBD1 mRNA expression was measured for IMU (154% ± 33 at 1:100 dilution) and extract SINx (144% ± 27 at 1.67mg/ml) as compared to control (figure 3). A similar trend was observed in LL-37 regulation upon co-treatment with IMU (184%±68 at 1:60 dilution), BRO-TP (161% ± 17 at 0.167 mg/ml) or BRO-TE (130% ± 2 at 1:400 dilution) together with the SARS-CoV-2 spike protein. Additionally, the extract-induced secretion of HBD1 and LL37 was analysed by ELISA in presence and absence of the spike protein at different time points (figure 4). Both extracts, IMU and BRO TP, could significantly trigger also the secretion of HBD1 peptide with or without S1 spike protein co-treatment from A549 cells. Extract BRO TP could also significantly trigger the secretion of LL-37 either with S1 spike protein co-treatment or without. This was not the case for extract IMU.

**Figure 3:**
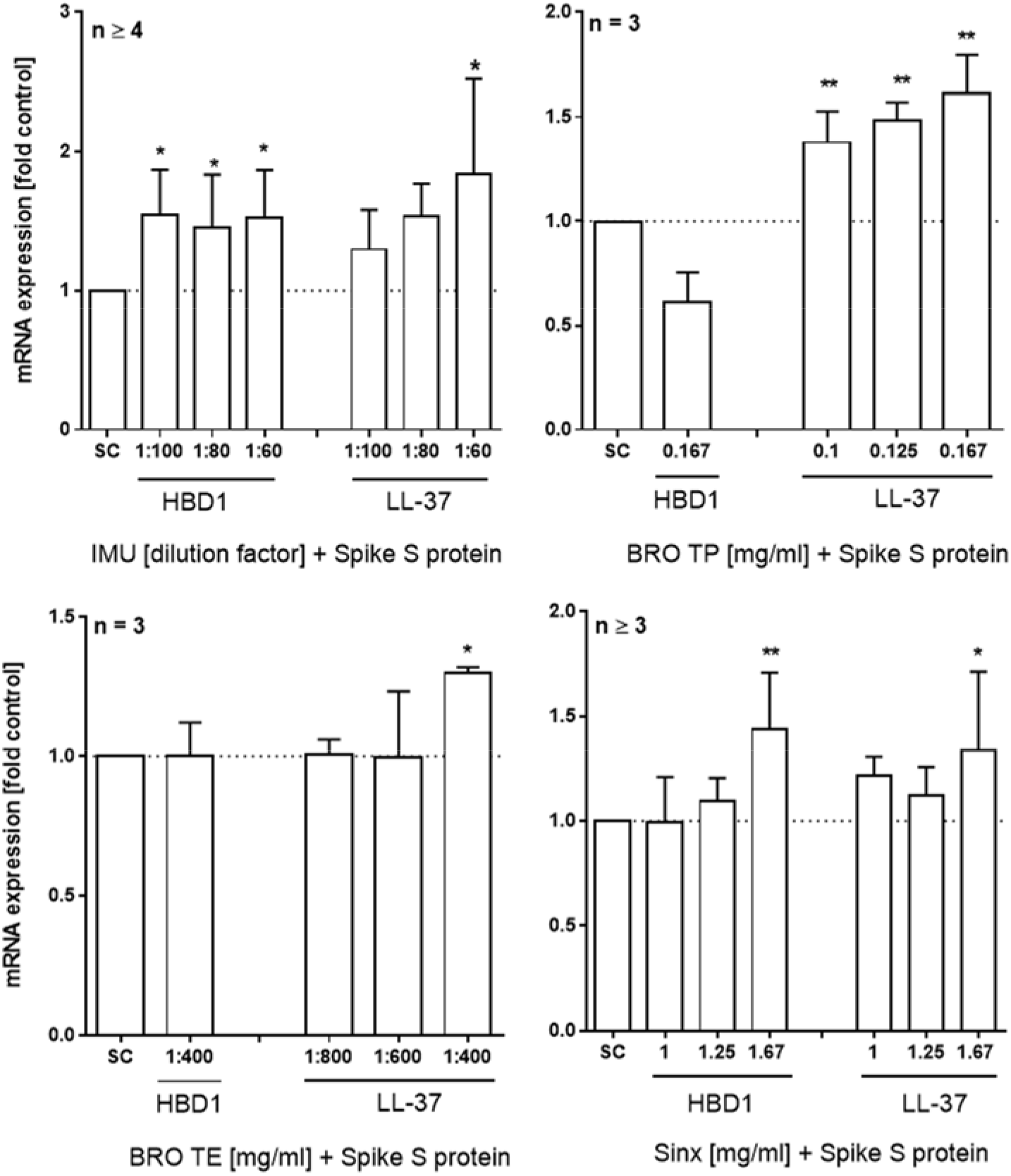
Effect of extracts on HBD1 and LL-37 mRNA expression in Spike protein stimulated A549 lung cells. A549 cells were cultured for 25 days and co-treated with extracts and SARS-CoV-2 Spike S1 protein for 24h. Bars are mean values +SD of at least three independent experiments.

**Figure 4:**
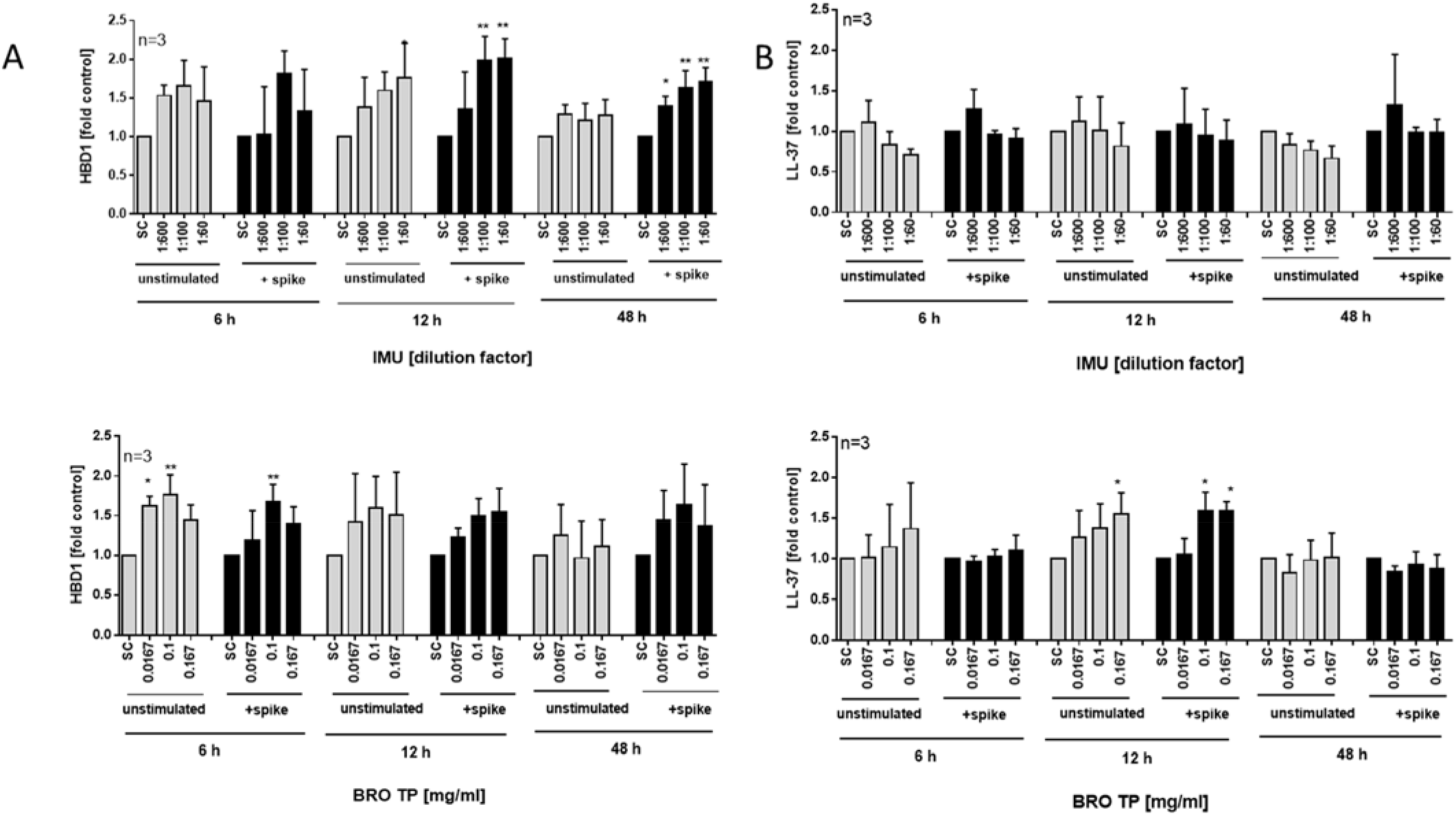
Effects of extracts on HBD1 and LL-37 peptide section measured by ELISA technique. Human A549 lung cells were cultured for 25 days and exposed to A) IMU or B) BRO TP for different time points with and without Sars-CoV-2 Spike S1 protein stimulation. Bars are mean values +SD of at least three independent experiments.

## Discussion

Herbal extracts are known to induce diverse cellular defense mechanisms following viral infection of human cells (Khare et al., 2020; Kim et al., 2010). Using different screening assays, we show here that extracts of marketed herbal medicinal products elicit potential beneficial effects *in vitro* in terms of cellular defense activation upon challenge with SARS-CoV-2 spike protein or peptide mix.

The first step of viral infection is the interaction of the virus with the host cell. In case of SARS– CoV-2 the spike protein interacts with the ACE2 receptor on the surface of epithelial cells such as from in the oral cavity or respiratory tract (Mani et al., 2020; Tay, 2020; Xu et al., 2020). In our study, extracts from IMU and BRO TE were able to interfere with the binding between the S1 spike protein and the human ACE2 receptor. Computational binding studies show that common herbal secondary metabolites like luteolin or quercetin could be able to bind and block the ACE2 receptor (Khare et al., 2020; Mani et al., 2020; Muchtaridi et al., 2020), and also bind the SARS-CoV-2 spike protein. A demonstration of interference with SARS-CoV-2 spike –ACE2 binding using cell-free or *in vitro* assays is missing for most of them, though. So, for quercetin and its metabolites, inhibition of recombinant human ACE2 (rhACE2) activity has been reported *in vitro* (Liu et al., 2020). The rhACE2 activity was then inhibited by rutin, quercetin-3-O-glucoside, tamarixetin, and 3,4-dihydroxyphenylacetic acid by 42–48%. With an IC50 of 4.48 μM, quercetin was the most potent rhACE2 inhibitor tested in this study. The herbal extracts investigated here contain high amounts of these plant metabolites (Assessment report on Quercus robur L.; EMA, 2013; EMA/HMPC/136583/2012 Assessment report on Primula veris L. and/or Primula elatior (L.) Hill, 2012; EMA/HMPC/342334/2013 Assessment report on Thymus vulgaris L. and L., 2012; Seibel et al., 2015; Seibel et al., 2018). Thus, these constituents could account or add to the inhibitory effect observed by the two herbal extract mixtures IMU and BRO TE and it will be important to further ascertain this hypothesis in ongoing studies. The products are mixtures made from several medicinal plants, thus it might also be helpful to determine the plant with the largest share on this effect. This is the more important as the inhibition was observed at a concentration range which turned out to be cytotoxic to human PBMC and which is not relevant for the direct use in practice. But, IMU and BRO TP also activated the human innate immune defense by increasing the level of defensin HBD1 and/or cathelicidin LL37 upon SARS-CoV-2 spike protein stimulation and this was evident at much lower concentrations. By regulating chemokine and cytokine production, both AMPs help to maintain homeostasis of the immune system and display antiviral properties, as for example evidenced by gene regulation upon viral challenge or expression in cells involved in viral defense (Wilson et al., 2013). LL-37, consisting of 37 amino acids and an overall positive net charge (+6) can also eliminate microbes directly by electrostatic binding to negatively charged molecules on microbial membranes (PrePrint Roth et al., 2020). In secretions from lung and nose it was found that LL-37 could reach high concentrations indicating to a relevant role in lung immune defense mechanisms (Kim et al., 2003; Mansbach et al., 2017; Schaller-Bals et al., 2002). Defensins can be detected in the mucosa of all respiratory tissues, including pharynx (22, 23). The effect of AMPs on virus infections appear to be specific to the virus, AMP, and also target cell and to our knowledge, a direct antiviral efficacy of HBD-1 or LL-37 against SARS-CoV-2 has not been shown so far (Ding et al., 2009). However, interestingly, for LL-37 a binding to the SARS-CoV-2 spike protein and an inhibitory action of LL-37 on the spike protein to its entry receptor have been reported using binding competition studies (PrePrint Roth et al., 2020). In another study, high structural similarity of LL-37 to the N-terminal helix of the receptor-binding domain of SARS-CoV-2 was reported (PrePrint Kiran Bharat Lokhande, 2020). There is also currently speculation that vitamin D, by upregulation of LL-37, could be beneficial on the course of COVID-19 disease (Crane-Godreau et al., 2020) and LL-37 even has been proposed by some researchers for treatment of COVD-19 patients (PrePrint Zhang et al., 2020). Thus, stimulating LL37 expression, as observed here *in vitro* with BRO TP, might have several advantages during early phase of SARS-CoV-2 entry, but this hypothesis, again, needs further verification.

BRO TE and SINx extracts could be shown in the present study to further boost the immune response of PBMC in terms of intracellular IFN-γ expression, which was activated by a mixture including SARS-CoV-2 spike peptides. Once a virus has entered the cell and is replicated, a host immune response starts to combat viral infection (Tay, 2020). An early interferon response of the host has been reported to be essential for an effective defense against SARS-CoV-2 infection (Liao et al., 2020; Tay, 2020). In turn, a decrease of IFNγ positive T-helper cells might increase the risk for severe courses of COVID-19 (Pierce et al., 2020; Sattler et al., 2020). The here reported *in vitro* findings provide a first hint that BRO TE and SINx may help in further IFNγ expression during infection. On the one hand, INF-γ is essential for the antiviral defense, but on the other hand persistant high levels of INF-γ have been reported to worsen the systemic inflammation, intensifying tissue injury and organ failure (Wang et al., 2020) during COVID-19 disease. This yet known ambiguous role of IFN-γ in the course of SARS-CoV-2 infection here also asks for special attention.

In conclusion, the marketed herbal medicinal products tested in this study demonstrate a range of potential support of the human immune defense against SARS-CoV-2 infection. Whether the observed effects could be relevant for the systemic use in man or limited to local effects e. g. in the oral cavity, needs to be investigated. Further confirmatory mechanistic studies are necessary to gain a deeper understanding of the reported observations and the transferability of the *in vitro* results to the clinical situation needs to be tested.

## List of Abbreviations

|  |  |
| --- | --- |
| ACE2 | Angiotensin-converting enzyme 2 |
| AMP | Antimicrobial peptide |
| BRO-TE | Bronchipret® thyme-ivy |
| BRO-TP | Bronchipret® thyme-primrose |
| CC<sub>50</sub> | Cytotoxic concentration |
| COVID-19 | Coronavirus disease 2019 |
| DER | Drug-extract ratio |
| IMU | Imupret® |
| PMA | Phorbol-12-myristat-13-acetat |
| PBS | Phosphate-buffered saline |
| FCS | Fetal calf serum |
| qRT-PCR | Quantitative reverse transcriptase polymerase chain reaction |
| SARS-CoV-2 | Severe acute respiratory syndrome coronavirus 2 |
| SD | Standard Deviation |
| SINx | Sinupret extract® |
| TOP | Tonsipret® |

## Declarations

### Ethics approval and consent to participate

The study on human PBMC was approved by the Ethics Committee of the University of Freiburg and carried out according to their guidelines and regulations (ethical vote 373/20).

### Consent for publication

Not applicable. This article does not contain data from any individual person.

### Availability of data and material

The datasets used and/or analyzed during the current study are available from the corresponding author on request.

### Authors’ contributions

Study design: E.L‥All authors contributed to the study conception, Design of experiments, data acquisition, analysis of data: H.T.T, E. L. The first draft of the manuscript was written by H.T.T. and E.L. All authors commented on previous versions of the manuscript.

